# Midbrain and lateral nucleus accumbens dopamine depletion affects free-choice high-fat high-sugar diet preference in male rats

**DOI:** 10.1101/2020.11.17.384651

**Authors:** Anil Joshi, Fanny Faivre, Susanne Eva la Fleur, Michel Barrot

## Abstract

Dopamine influences food intake behavior. Reciprocally, food intake, especially of palatable dietary items, can modulate dopamine-related brain circuitries. Among these reciprocal impacts, it has been observed that an increased intake of dietary fat results in blunted dopamine signaling and, to compensate this lowered dopamine function, caloric intake may subsequently increase. To determine how dopamine regulates food preference we did 6-hydroxydopamine (6-OHDA) lesions, depleting dopamine in specific brain regions in male Sprague Dawley rats. The food preference was assessed by providing the rats with free choice access to control diet, fat, 20% sucrose and tap water. Rats with midbrain lesions targeting the substantia nigra (which is also a model of Parkinson’s disease) consumed fewer calories, as reflected by a decrease in control diet intake, but they surprisingly displayed an increase in fat intake, without change in the sucrose solution intake compared to sham animals. To determine which of the midbrain dopamine projections may contribute to this effect, we next compared the impact of 6-OHDA lesions of terminal fields, targeting the dorsal striatum, the lateral nucleus accumbens and the medial nucleus accumbens. We found that 6-OHDA lesion of the lateral nucleus accumbens, but not of the dorsal striatum or the medial nucleus accumbens, led to increased fat intake. These findings indicate a role for lateral nucleus accumbens dopamine in regulating food preference, in particularly the intake of fat.

**HIGHLIGHTS:**

- Dopamine influences fat intake
- Animal model of Parkinson’s disease display lower kcal intake but increased fat choice
- Dopamine depletion in the lateral nucleus accumbens favors fat intake

## INTRODUCTION

Food intake is a highly reinforced behavior that not only provides nutrients needed for survival, but that also induces feelings of rejoice and pleasure (Hoebel, 1985). Modern human diet and natural rewards include a substantial amount of sugars and fats, which can modulate the dopamine signaling in the brain reward system (DiFeliceantonio et al., 2018; Fernandes et al., 2020; Fritz et al., 2018) and could disturb the internal homeostatic mechanism which regulates hunger and satiety, ultimately leading to overconsuming and obesity (de Araujo et al., 2008; Han et al., 2018; Zimmerman and Knight, 2020).

Homeostatic signals that influence the excitability of midbrain dopamine neurons can influence the sensitivity to drugs (Volkow et al., 2017) and it is an ongoing debate whether the palatable food addiction and drug addiction do share a similar mechanism (Fletcher and Kenny, 2018). Dopamine is one of the important neurotransmitters involved in reward processing, including in the rewarding aspects of food intake (Berke, 2018; Cox and Witten, 2019; Hernandez and Hoebel, 1988). The main sources of the dopamine system are dopamine cell bodies in the substantia nigra par compacta (SNc) and ventral tegmental area (VTA) that project to the striatal complex (Gerfen and Bolam, 2016). Classically, the VTA dopamine neurons that project to ventral striatum have been linked to reward processes (Hamid et al., 2016; Hu, 2016; Lammel et al., 2012; Morales and Margolis, 2017), whereas SNc dopamine neurons are more often associated with motor control (da Silva et al., 2018; Howe and Dombeck, 2016). However, this dichotomy has been challenged and several studies have shown that SNc neurons projecting to the dorsal striatum can also be linked to food-motivation (Ilango et al., 2014; Lee et al., 2020; London et al., 2018) and associated movements (Lee et al., 2020). Little is known however about SNc striatal projections in relation to obesogenic diets. Dopamine deficient mice lack goal-directed feeding behavior and die of starvation by the age of 4 weeks (Palmiter, 2008); however, if the dopamine signaling is restored in the striatum (Szczypka et al., 2001) or in the DA neurons that project to the dorsal striatum (Hnasko et al., 2006; Sotak et al., 2005), then these mice consume enough for survival. Similarly, feeding inhibition is seen when dopamine signaling is altered in the dorsal striatum (Sotak et al., 2005).

As observed for drugs of abuse, palatable food can elicit altered dopaminergic signaling in the striatum. It has, for example, been shown that intragastrical fat infusions increase extracellular dopamine levels in the dorsal striatum (Ferreira et al., 2012). Dopamine signaling is also altered in the nucleus accumbens during the consumption of diets that are rich in fat and/or sugar (Hajnal et al., 2004; Liang et al., 2006; Rada et al., 2012) or during food restriction (Carr, 2020). An excessive consumption of fats has been shown to reduce brain dopaminergic function, and it has been hypothesized that this dopamine deficiency may aggravate obesity by favoring overfeeding as a compensatory mechanism to restore reward sensitivity (Tellez et al., 2013). A palatable high-fat diet has thus been proposed to promote addictive-like behavior in rats by downregulating D_2_ receptors while promoting obesity (Johnson and Kenny, 2010), which is in agreement with human PET scanning in obese subjects showing lower availability of D_2_ receptor in the striatum (de Weijer et al., 2011; Wang et al., 2001). It has also been shown that rats under a free-choice high-fat high-sugar (fcHFHS) paradigm have decreased striatal D_2_ receptor availability (van de Giessen et al., 2013), and that male rats fed with a diet rich in saturated dietary lipids (palm oil) have reduced D_1_-mediated signaling in the nucleus accumbens (Hryhorczuk et al., 2016). However, the influence that dopamine projections reciprocally exert on the overconsumption of fat and/or sugar remains to be detailed.

In the present study, we used 6-hydroxydopamine (6-OHDA) lesions of midbrain dopamine neurons and of specific projections within the striatal complex to study the influence of dopamine in the fcHFHS paradigm, an obesogenic diet known to facilitate obesity and insulin resistance (la Fleur et al., 2011; la Fleur et al., 2010). This study, conducted in the rat, highlights a role for dopamine in the lateral nucleus accumbens in fat choice and intake.

## EXPERIMENTAL PROCEDURES

### Animals

Adult male Sprague-Dawley rats (Janvier Labs, France) were housed under standard conditions (2 per cage, 22±1°C, 12-h light-dark cycle) and habituated to the animal facility for at least a week before starting procedures. A total of 128 rats were purchased for the various experiments, and 118 rats of them were finally used for food intake behavior. Animal care and use were performed in accordance with the Centre National de la Recherche Scientifique (CNRS) and the European Union directives, with procedures approved by the regional ethical committee (CREMEAS).

### Surgical procedures

Rats (7-8 weeks old; 250-325g) were anesthetized using either ketamine (Imalgene® 1000) / xylazine (Rompun® 2%) / acepromazine (Calmivet®) (80 mg/kg, 4 mg/kg and 1 mg/mL respectively; intraperitoneal (i.p.)) or using zolazepam (Zoletil® 50) / xylazine (Rompun®) (65 mg/kg and 14 mg/kg respectively; i.p.), and placed in a stereotaxic frame (Kopf Instruments). Ocry-gel® was used to protect the eyes; and subcutaneous bupivacaine (40 µL/100 g) and Metacam® (50 µL/100 g) were used as local anesthetic for surgery and as anti-inflammatory drug for postsurgical pain relief respectively. Stereotaxic coordinates relative to the bregma were adjusted to the animal weight based on trial animals with blue dye injections. Coordinates (in mm) were as follows (Paxinos and Watson, 2013): SNc, anteroposterior (AP)=-5.1, lateral (L)=±2.2, vertical (V)=-7.4; dorsolateral striatum, AP=+1.7, L=±3.1, V=-4.2; medial nucleus accumbens (mNAc), AP=+1.8, L=+1.5, V=-6.9; and lateral nucleus accumbens (lNAc), AP=1.5, L=±2.6, V=-7.0. Verticality was taken from the dura. Hamilton syringes with 33 gauge needles were used to deliver 2 µL of 6-OHDA (Sigma-Aldrich, France; 2.5 µg/µL in 0.9% NaCl with 0.01% ascorbic acid) (Faivre et al., 2020) bilaterally, over 4 minutes. Needles were removed 7 minutes after the end of the injection. The sham surgeries were performed in a similar way, without injection. Rats were allowed to recover for 3 weeks before starting the behavioral experiments.

### Free-choice High-Fat High-Sugar experiments

The fcHFHS paradigm (la Fleur et al., 2014) was implemented to assess food/fluid choice and intake following 6-OHDA lesions. The test was performed over 7 days. Rats were single housed and given free excess to normal chow (diet A04, SAFE, France), fat (Les Pâturages des Flandres pure graisse de bœuf, Sarl Buchez, estaires, France), 20% sucrose solution (Sigma-Aldrich), and water. On each day, the position of the water and sucrose bottles was interchanged, with a change in fat and chow positions every other day. The SNc lesion experiment initially included 25 sham and 27 lesioned rats, the dorsolateral striatum lesion experiment 8 sham and 8 lesioned rats, the mNAc lesion experiment 10 sham and 14 lesioned rats, the lNAc lesion experiment 12 sham and 14 lesioned rats.

### Immunohistochemistry

After completion of behavioral testing, rats were killed with a sodium pentobarbital overdose (235 mg/kg, i.p.) and decapitated. Their brains were collected and fixed in a paraformaldehyde solution (4% in 0.1 M phosphate buffer (PB), pH 7.4) with 20% glycerol for 24 hours, and transferred to a 20% sucrose solution in PB for further cryoprotection. Coronal sections (40 μm) were cut using a cryotome (Leica SM 2000R). Immunohistochemistry against tyrosine hydroxylase (TH) was performed as previously described (Faivre et al., 2020). At room temperature, free-floating sections were washed 3 times with 0.9% NaCl / 0.01 M PB pH 7.4 (PBS), incubated in 50% ethanol with 0.03% hydrogen peroxide for 30 min for endoperoxidase blocking, washed 3 times with PBS, incubated in 0.3% Triton X-100 in PBS (PBS-T) with 5% normal donkey serum for permeabilization and blocking for 45 min, and incubated overnight with the anti-TH primary antibodies (Millipore-Chemicon MAB318, 1/5000 for SNc sections and 1/2000 for striatal/NAc sections) in PBS-T with 1% normal donkey serum. On the following day, sections were washed 3 times with PBS, incubated with an anti-IGg biotinylated secondary antibody (#BA2001, Vector Laboratories 1/200) in PBS for 90 min, washed 3 times with PBS, and incubated with the avidin–biotin–peroxidase complex (ABC Elite; Vector Laboratories). The amplified complex was finally revealed with a peroxidase / 3.3’-diaminobenzidine tetrahydrochloride reaction. The sections were mounted, dried with a gradient of alcohol followed by Roti®-Histol solution, and a glass coverslip was placed using Eukitt®.

### Microscopy and analysis

The mounted slides were scanned in a NanoZoomer S60 Digital slide scanner C13210 (Hamamatsu) at 20x in bright field. Images were saved in Nanozoomer Digital Pathology Image (ndpi) format and the uncompressed images were viewed in NDP.view 2.6.13 and extracted in .tif format for analyses using Image J. The images were converted into greyscale and the grey densities were calculated on 5-6 sections per animal and per region. Striatum and NAc data were expressed as optical density, and SNc/VTA data were expressed as % of the mean integrated grey density in controls. Sections ranging from AP +2.2 to +1.2 from bregma were selected for analyses of the striatum and NAc, while sections ranging from AP - 4.6 to -5.8 were used for SNc and VTA analyses.

### Statistics

Results are expressed as mean±SEM in figures. Statistical analyses were performed using STATISTICA 13 software (Statsoft, Tulsa, OK, USA) and GraphPad Prism 8.04 (GraphPad Software, La Jolla California USA). Student’s t-test was used for two group comparison for lesion extent and for total kcal intake. ANOVA was used for comparing different food/fluid intakes over days (within factors) between the sham and lesion groups (between factor), followed by Duncan *post hoc*, with the level of significance at *p* < 0.05.

## RESULTS

### Effects of substantia nigra lesion on High-fat High-sugar free choice feeding

The SNc was lesioned using 6-OHDA and the extent of the lesions at cell body and terminal levels was controlled by TH immunohistochemistry at the end of the experiments (Fig. 1). Lesioned animals were considered for the analysis of behavioral data if they bilaterally displayed at least a 50% loss in SNc TH staining (Fig. 1F), which concerned 22 out of the 27 rats that received 6-OHDA. In those animals, the lesions concerned the SNc, but could also extend to the lateral parts of the VTA (Fig. 1E).

**Figure 1:**
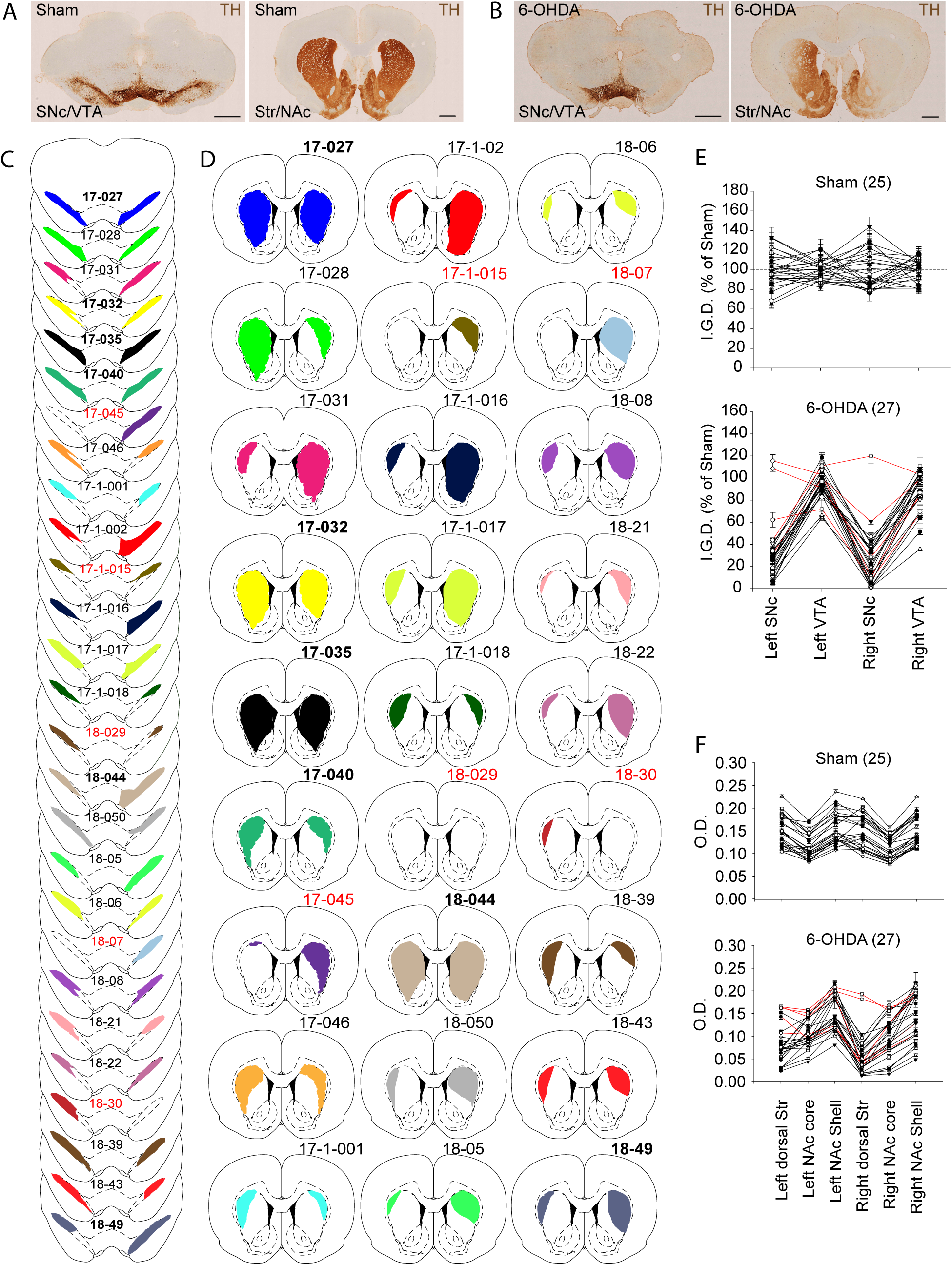
Midbrain 6-OHDA lesions. Examples of tyrosine hydroxylase (TH) immunostaining in the SNc/VTA area and in the Str/NAc area for Sham (**A**) and for 6-OHDA (**B**) injected rats (scale bars, 1mm). Individual extent of the lesion: the colored area corresponds to the lesioned area at the site of injection (**C**) and at Str/NAc projection level (**D**) after 6-OHDA injection. The experimental identity number of each rat is displayed nearby drawings; the red color identifies animals with less than 50% of bilateral TH immunostaining loss in the SNc that were removed from behavioral analyses. (**E**) Graph representing the integrated grey density (I.G.D.) of TH immunostaining in the SNc and VTA. (**F**) Optical density (O.D.) of TH immunostaining in different subregions of the Str/NAc. The number of animals is given between brackets. 6-OHDA, 6-hydroxydopamine; NAc, nucleus accumbens; SNc, substantia nigra pars compacta; Str, striatum; VTA, ventral tegmental area.

The fcHFHS paradigm was conducted over 7 days. Overall, the SNc lesion led to a decrease in total kcal intake *per* 100 g body weight (F_1,45_ = 4.41, *p* = 0.04) (Fig. 2A). With time, the rats also displayed slightly lower body weight than control (F_4,200_ = 7.29, *p* < 0.0001) (Supplementary Fig. 1S). Interestingly, the decrease in kcal intake was associated with a change in the nutrient choice (F_2,90_ = 8.23, *p* = 0.0004): the rats decreased their chow intake (post-hoc, *p* = 0.00002), increased their fat intake (*p* = 0.01), and maintained their sugar intake (*p* = 0.44). As total intake differed between lesioned animals and their controls, we also standardized the data (% of intake for each animal) (Fig. 2B), which confirmed the increased intake of high-fat food to the detriment of regular chow (F_2,90_ = 4.88, *p* = 0.009; *post hoc*: decreased chow *p* = 0.006, increased high-fat *p* = 0.006, unchanged sucrose *p* = 0.99). This difference appeared as robust as it was stable over days (F_12,540_ = 1.42, *p* = 0.15).

**Figure 2:**
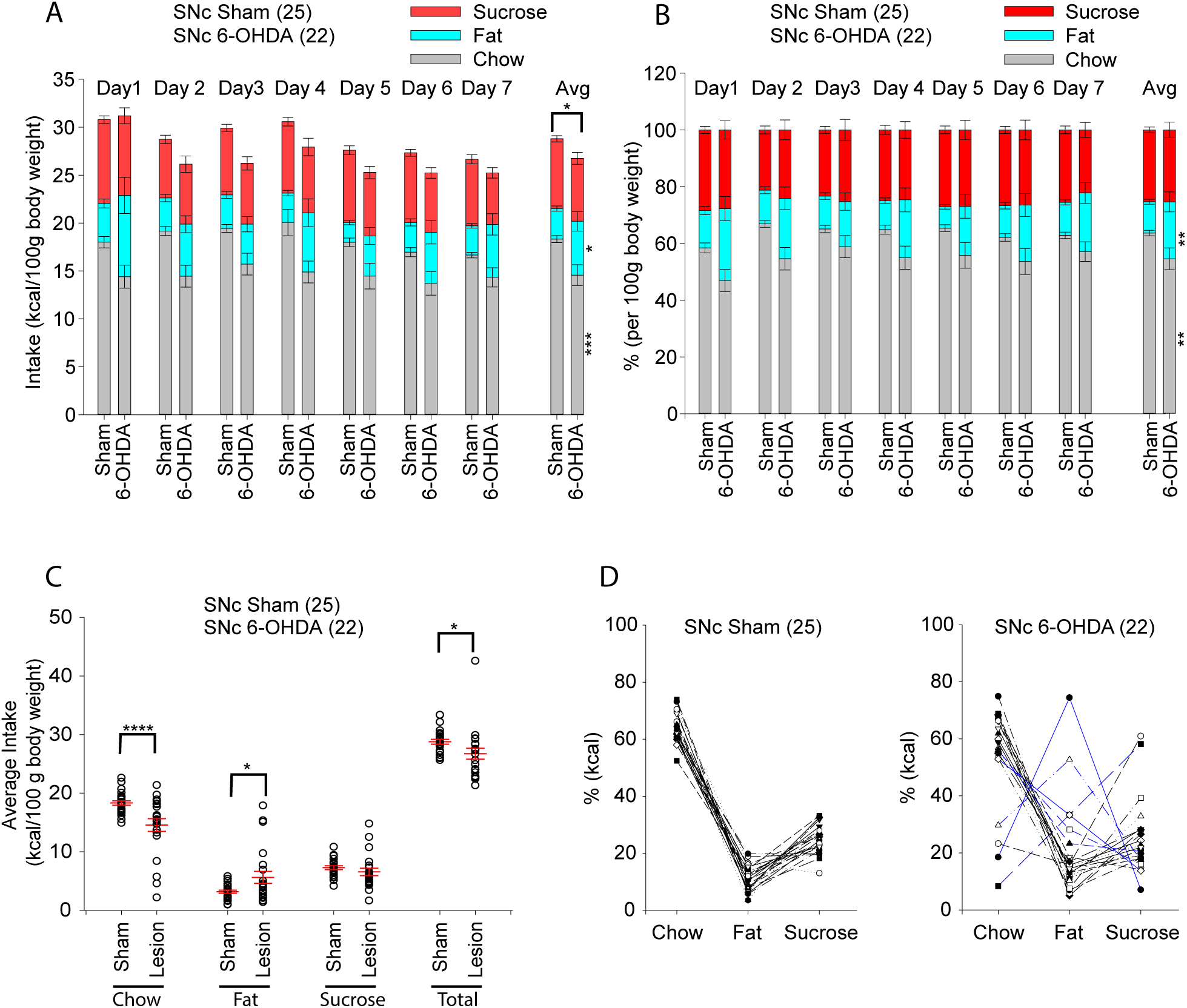
High-fat High-sugar free choice in SNc lesioned animals. Daily nutrient intake pattern in sham and 6-OHDA animals for the three nutrient types over 7 days, in kcal per 100 gram of body weight (**A**), and in percentage of kcal consumed from different nutrient types (**B**). (**C**) Individual average kcal intake per 100 g of body weight in sham and 6-OHDA animals. (**D**) Individual patterns of nutrient choice (% of kcal intake) in sham and 6-OHDA animals. The number of animals is given between brackets. Data are presented as mean±SEM. *, p < 0.05; **, p < 0.01; ***, p < 0.001; ****, p <0.0001. 6-OHDA, 6-hydroxydopamine; Avg, average; SNc, substantia nigra pars compacta.

When looking more closely at the individual pattern of nutrient choice (Fig. 2D; Supplementary Fig. S2), a highly stable pattern was observed among the 25 Sham animals. This pattern of choice was however strongly disrupted in some of the 6-OHDA lesioned animals. In particular, the mean increase in high-fat food intake was in fact due to 6 animals only (27% of the cohort), and those animals were among the ones with the largest extent of striatal loss in TH staining (bold animal numbers in Fig. 1; blue lines in Fig. 2D). This observation led us to study more closely the role of dopamine terminals from different striatal subregions in the fcHFHS paradigm and high-fat food intake.

### Effects of dopamine depletion in the dorsolateral striatum on High-fat High-sugar free choice feeding

We first tested the influence of dopamine terminals in the dorsolateral striatum, by performing local 6-OHDA injections (Fig. 3). TH staining was used to control the extent of these lesions, which were limited to the lateral and dorsolateral parts of the dorsal striatum (Fig. 3C). With the fcHFHS paradigm, we observed no impact of the lesions on the total kcal intake *per* 100 g body weight (F_1,14_ = 0.05, *p* = 0.81) (Fig. 3D), or on the choice between regular chow, high-fat diet and sucrose solution (F_2,28_ = 1.38, *p* = 0.26) (Fig. 3E). The individual patterns of nutrient choice (Fig. 3G) further supported this lack of impact of the dorsolateral striatum lesions, except for one animal with higher fat intake (blue line in Fig. 3G) that displayed a ventrolateral extent of the lesion (picture insert in Fig. 3G). We then lesioned the dopamine terminals in the lNAc to test their influence on the fcHFHS paradigm and high-fat food intake.

**Figure 3:**
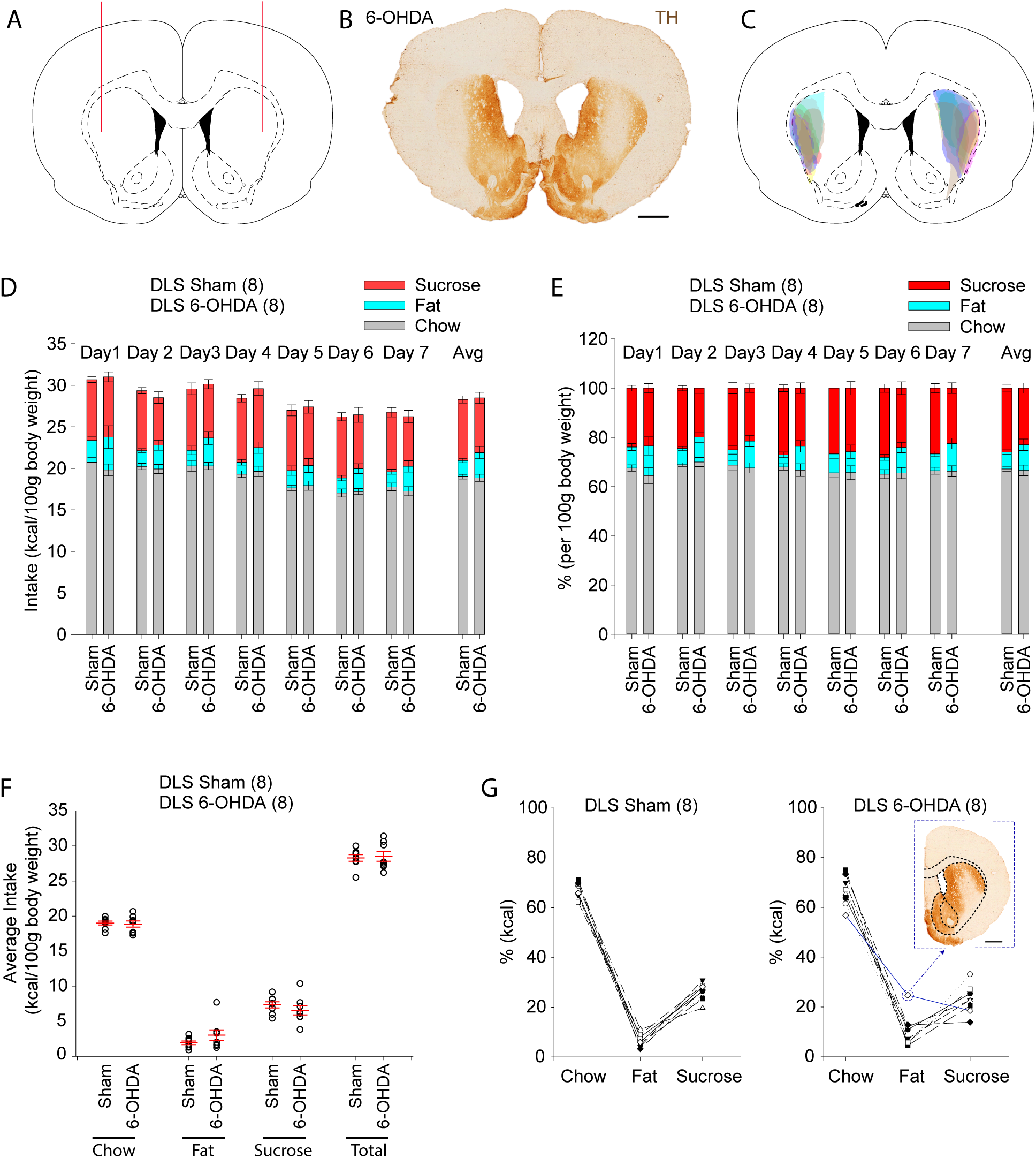
High-fat High-sugar free choice in DLS lesioned animals. Schematic of injection site (**A**), example of tyrosine hydroxy-lase immunostaining (**B**) and schematic reconstruction of individual lesion extents (**C**). Daily nutrient intake pattern in sham and 6-OHDA animals for the three nutrient types over 7 days, in kcal per 100 gram of body weight (**D**), and in percentage of kcal consumed from different nutrient types (**E**). (**F**) Individual average kcal intake per 100 g of body weight in sham and 6-OHDA animals. (**G**) Individual patterns of nutrient choice (% of kcal intake) in sham and 6-OHDA animals. The picture insert displays TH immunostaining in the animal with high fat intake. The number of animals is given between brackets. Data are presented as mean±SEM. Scale bars, 1 mm. 6-OHDA, 6-hydroxydopamine; Avg, average; DLS, dorsolateral striatum.

### Effects of dopamine depletion in the nucleus accumbens on High-fat High-sugar free choice feeding

The 6-OHDA lesion of the lNAc (Fig. 4) led to an overall increase in total kcal intake *per* 100 g body weight (F_1,24_ = 11.53, *p* = 0.002) (Fig. 4D). This increase was more specifically due to a change in the nutrient choice (F_2,48_ = 6.92, *p* = 0.002), with an increased intake of high-fat food (*post hoc, p* = 0.00006) and no quantitative change in chow (*p* = 0.72) and sucrose (*p* = 0.64) intake. When data were standardized (% of intake for each animal) (Fig. 4E), it confirmed the increased choice of high-fat food, mostly to the detriment of regular chow (F_2,48_ = 7.80, *p* = 0.001; *post hoc*: decreased chow *p* = 0.01, increased high-fat *p* = 0.0003, unchanged sucrose *p* = 0.18). The individual patterns of nutrient choice (Fig. 4G; Supplementary Fig. S2) further supported the overall increase in high-fat food intake.

**Figure 4:**
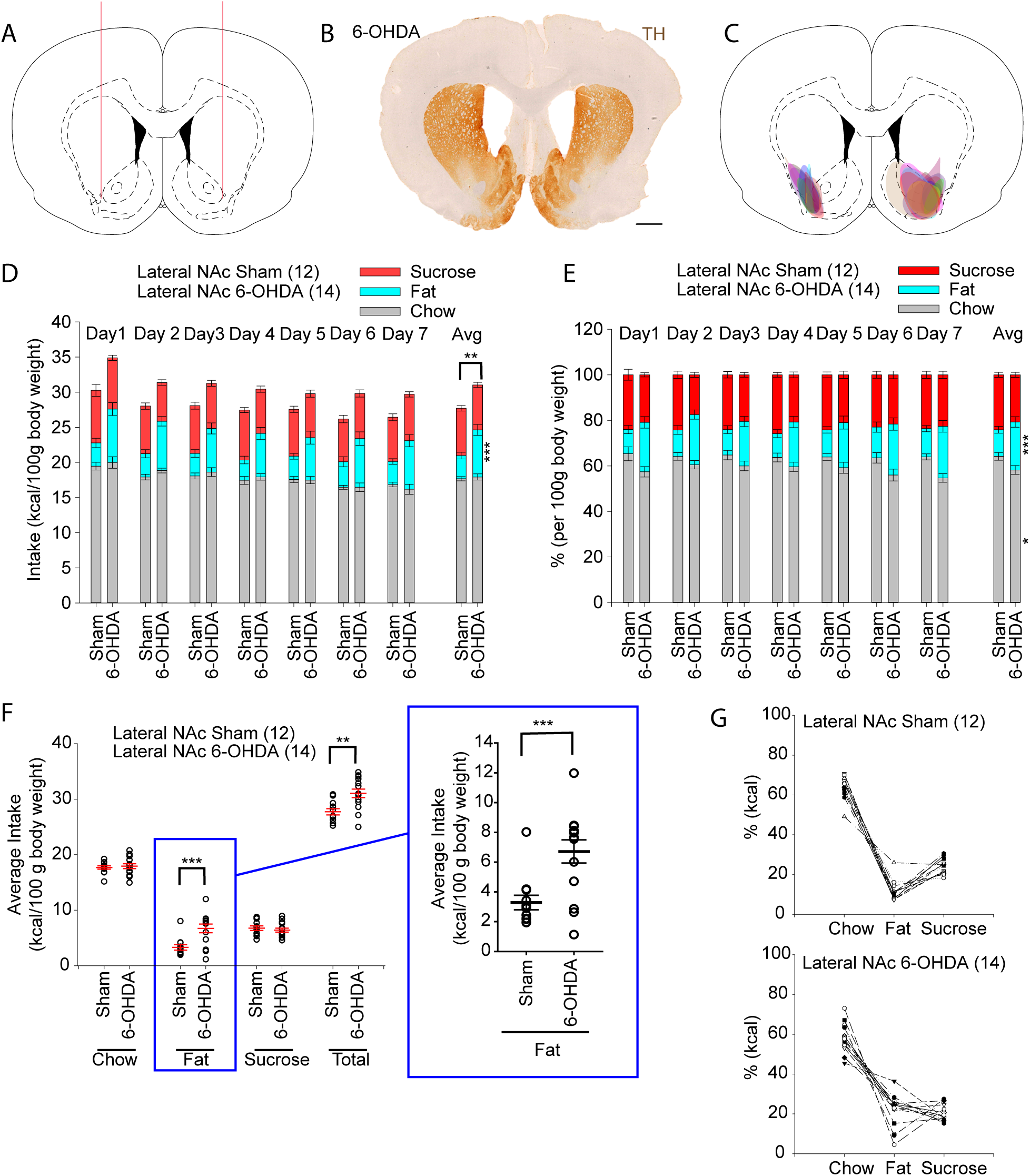
High-fat High-sugar free choice in lateral NAc lesioned animals. Schematic of injection site (**A**), example of tyrosine hydroxylase immunostaining (**B**) and schematic reconstruction of individual lesion extents (**C**). Daily nutrient intake pattern in sham and 6-OHDA animals for the three nutrient types over 7 days, in kcal per 100 gram of body weight (**D**), and in percentage of kcal consumed from different nutrient types (**E**). (**F**) Individual average kcal intake per 100 g of body weight in sham and 6-OHDA animals, the insert details the individual fat intake. (**G**) Individual patterns of nutrient choice (% of kcal intake) in sham and 6-OHDA animals. The number of animals is given between brackets. Data are presented as mean±SEM. *, p < 0.05; **, p < 0.01; ***, p < 0.001. Scale bar, 1 mm. 6-OHDA, 6-hydroxydopamine; Avg, average; NAc, nucleus accumbens.

We then tested whether this role could be extended to the rest of the NAc or was specific to the lNAc subregion. In the fcHFHS paradigm, the 6-OHDA lesion of the mNAc (Fig. 5) had no significant impact on the total kcal intake *per* 100 g body weight (F_1,22_ = 0.002, *p* = 0.96) (Fig. 5D), or on the choice between regular chow, high-fat diet and sucrose solution (intake: F_2,44_ = 0.50, *p* = 0.60; %: F_2,44_ = 0.44, *p* = 0.64) (Fig. 5D,E), which supported a specific influence of the lNAc on high-fat choice.

**Figure 5:**
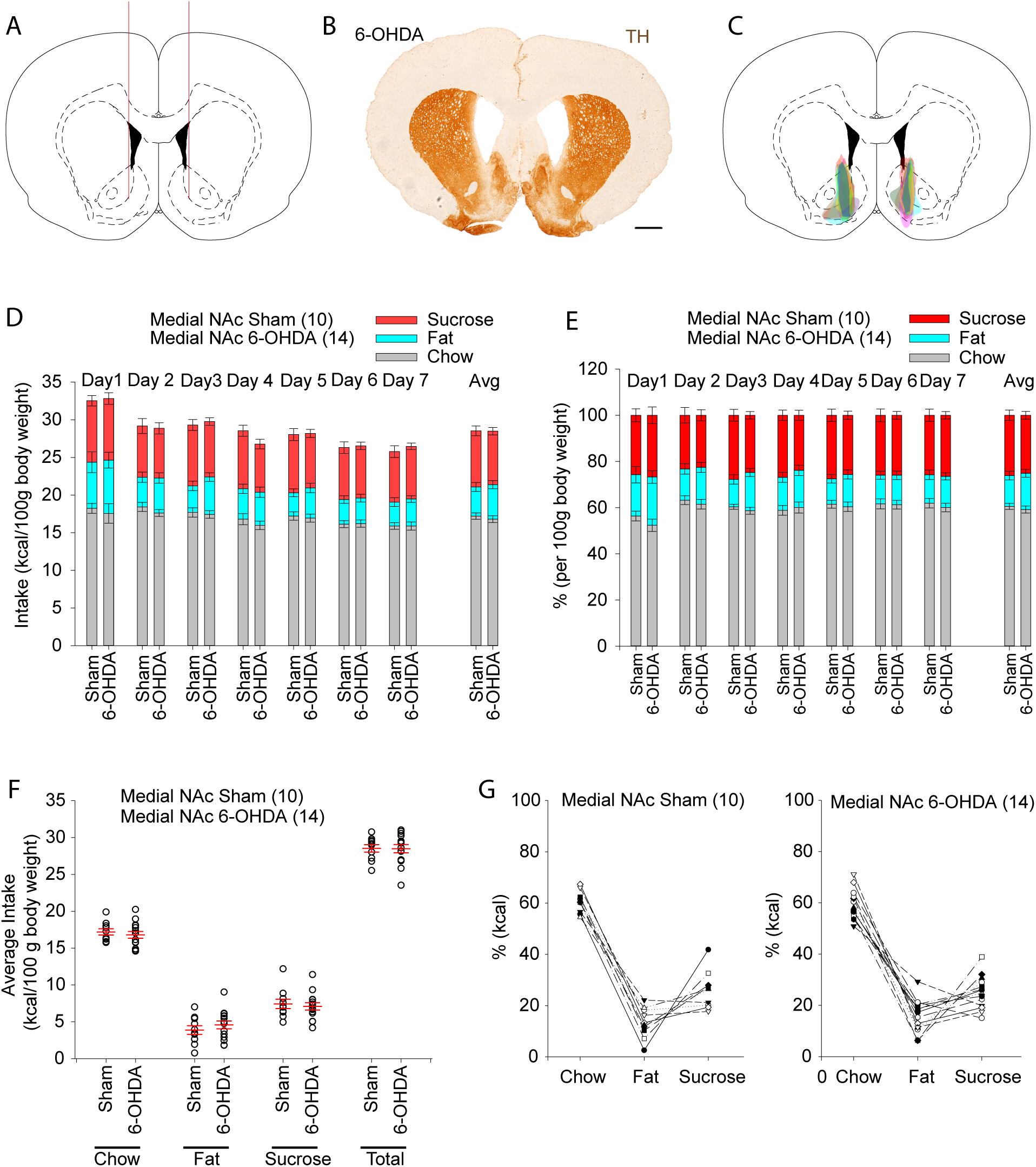
High-fat High-sugar free choice in medial NAc lesioned animals. Schematic of injection site (**A**), example of tyrosine hydroxylase immunostaining (**B**) and schematic reconstruction of individual lesion extents (**C**). Daily nutrient intake pattern in sham and 6-OHDA animals for the three nutrient types over 7 days, in kcal per 100 gram of body weight (**D**), and in percentage of kcal consumed from different nutrient types (**E**). (**F**) Individual average kcal intake per 100 g of body weight in sham and 6-OHDA animals. (**G**) Individual patterns of nutrient choice (% of kcal intake) in sham and 6-OHDA animals. The number of animals is given between brackets. Data are presented as mean±SEM. Scale bar, 1 mm. 6-OHDA, 6-hydroxydopamine; Avg, average; NAc, nucleus accumbens.

## DISCUSSION

In the present study, we used a free-choice high-fat high-sugar diet preference paradigm in rats in order to identify the impact of dopamine loss on food intake and preference. Our results highlighted a role of dopamine terminals in the lateral nucleus accumbens in the regulation of fat intake.

6-OHDA is a neurotoxin widely used to model Parkinson’s disease (Faivre et al., 2019) and to study reduced dopamine states. This toxin produces reactive oxygen species and causes the death of the catecholaminergic neurons and their terminals (Ungerstedt, 1968). The nigrostriatal dopamine system is important for feeding behavior as complete bilateral degeneration of SNc with 6-OHDA causes adipsia and aphagia (Ungerstedt, 1971). Similarly, dopamine deficient mice are hypoactive and aphagic, leading to starvation and death if they are not given L-Dopa (Palmiter, 2008). However, an appropriate post-operative care allows animals with partial bilateral SNc lesion to consume sufficient food and prevents animal loss (Faivre et al., 2020). In the present study, tests were performed after 3 weeks of post-surgical recovery, thus ensuring that animals were not having major problems of food and water intake.

As previously observed (Faivre et al., 2020), the animals with bilateral SNc lesion displayed lower body weight gain. This decreased weight gain may likely be due to the lower kcal intake from standard chow in comparison with sham animals. This reduced weight gain was however only observed with midbrain SNc lesions, but was not present with more selective terminal lesions. Decreased weight gain or even weight loss is not limited to the 6-OHDA model of Parkinson’s disease. Indeed, weight loss is also present with aging in the MitoPark transgenic mouse model (Li et al., 2013). Besides, mice expressing the A53T human alpha-synuclein mutation (Giasson et al., 2002), the Thy1-aSYN Mice (Cuvelier et al., 2018), and the PINK1 deficient mice (Gispert et al., 2009), also display reduced body weight gain over time. Similarly, in the MPTP (1-methyl-4-phenyl-1,2,3,6-tetrahydropyridine) model of Parkinson’s disease in the primate, a daily care of the animals is essential otherwise animals may die due to adipsia, aphagia and loss of body weight (Porras et al., 2012). However, it is to be noted that MPTP-treated mice (Munoz-Manchado et al., 2016; Sundstrom et al., 1990; Zhang et al., 2017) and the LRRK2 G2019S transgenic mice (Bieri et al., 2019) do not display noticeable change in body weight. Clinically, Parkinson’s disease patients are known to have weight fluctuations. Weight loss has been reported in early as well as advanced forms of the Parkinson’s disease (Kistner et al., 2014; Ma et al., 2018), whereas some patients show again increased weight gain only after dopamine replacement therapy, pallidotomy or subthalamic nucleus deep brain stimulation (Kistner et al., 2014; Ma et al., 2018). In the animals, food choice paradigms were however not yet applied to the 6-OHDA SNc lesion model. The use of the fcHFHS paradigm allowed us to observe that, beside the expected decreased in kcal intake, SNc lesioned animals did not display notable change in sucrose consumption but had a perturbed food intake pattern, with about a third of the animals surprisingly displaying an increased kcal consumption from the fat source. This heterogeneity in individual data suggested that the subpopulation of lesioned dopamine neurons may be of importance and could explain interindividual variability in the consequences of the lesion. We thus targeted different dopamine terminal fields to test their influence on the fcHFHS paradigm.

The pharmacological or viral-mediated rescue of dopamine system in the dorsal striatum has been shown to restore feeding behavior in dopamine (tyrosine hydroxylase) deficient mice (Hnasko et al., 2006; Palmiter, 2008; Sotak et al., 2005; Szczypka et al., 2001; Zhou and Palmiter, 1995). These mice could indeed consume sufficient food for survival when a recombinant adeno-associated virus was delivered in the central or lateral region of the caudate putamen to ectopically co-express tyrosine hydroxylase and the GTP-cyclohydrolase (required by striatal cells to make tetrahydrobiopterin, an essential co-factor for tyrosine hydroxylase) (Szczypka et al., 2001), or when a retrograde canine adenovirus expressing the tyrosine hydroxylase was injected in the dorsal striatum (Sotak et al., 2005). Furthermore, a retrograde CAV2-Cre viral strategy could rescue feeding in tyrosine hydroxylase deficient mice by simply restoring the tyrosine hydroxylase gene selectively in neurons that project to the caudate putamen (Hnasko et al., 2006). These data converge to show that dopamine in the dorsal striatum may be critical to the aphagia associated with dopamine loss, and could partly explain the food-intake deficit that can be observed after bilateral SNc lesion. However, a 6-OHDA lesion that targets dopamine terminals only in the dorsolateral striatum is not sufficient *per se* to significantly alter weight gain, food and kcal intake, and preference for a specific food type. While this lack of impact was observed overall, some effect on food preference (increased fat intake) was however present in an animal whose lesion was extended to the ventrolateral part of the striatal complex.

Previous studies have shown that activation of VTA dopamine neurons projecting to the NAc is reinforcing (Tsai et al., 2009; Witten et al., 2011) and favors the frequency of feeding with smaller meal size, without affecting total food intake (Boekhoudt et al., 2017). When a D1 dopamine receptor agonist is given peripherally, it dose dependently affects palatable food consumption (Cooper et al., 1992; Martin-Iverson and Dourish, 1988). In this regard, the D1-expressing medium spiny neurons of the mNAC appears to be particularly sensitive to palatable food (Durst et al., 2019). Using recording and selective optogenetic manipulation of these neurons in the NAc shell, it has been shown that these neurons projecting to the lateral hypothalamus provide a rapid control over feeding and particularly feeding duration (O’Connor et al., 2015). Despite this well established and important control of fine aspects of feeding, we show that the more global overnight food intake and food choice is not altered by mNAc dopamine loss.

After high fat consumption, several studies have shown alterations in dopamine receptors, in dopamine transporter and in their mRNA levels in the nucleus accumbens (Adams et al., 2015; Hryhorczuk et al., 2016; Huang et al., 2005; Huang et al., 2006; Sharma and Fulton, 2013; South and Huang, 2008) and in the striatum (Alsio et al., 2010; Johnson and Kenny, 2010; Tellez et al., 2013). While fat intake can alter the dopamine system, less is reciprocally known on the role of the dopamine system in fat intake or preference. The binge consumption of sweetened high fat liquid in long Evans rats has for example been shown to be independent from NAc dopamine signaling (Lardeux et al., 2015), but this study was entirely carried out either in the core or medial NAc and did not address the potential influence of the lNAc.

Previously, it has been shown that bilateral 6-OHDA lesions of dopamine terminals in the NAc can enhance food intake during 30-min sessions (Koob et al., 1978). In the present study, we found that more specific bilateral lesions of the lNAc (but not of the mNAc) increased the daily total kcal intake, which was mainly driven by an increased intake of fat. This anatomical specificity within the NAc may be related to differences in connectivity within subregions of this nucleus. Indeed, the lNAc and the mNAc receive different dopamine innervation from the VTA, the lNAc receiving dopamine inputs from the lateral region of the VTA and projecting back in turn to GABAergic neurons of the lateral VTA causing disinhibition, while mNAc receives dopamine inputs from the medial region of the VTA and sends back GABAergic projection to both the medial and lateral VTA dopamine neurons thus causing inhibition (Yang et al., 2018). This organization may for example contribute to the information flow related to the ascending spiral moving from the ventral to the dorsal striatum (Haber et al., 2000). Furthermore, the VTA dopamine neurons projecting to the lNAc may receive afferents primarily from anterior cortical regions, including the prefrontal cortex, while the VTA dopamine neurons projecting to the mNAc may receive notable inputs from the dorsal raphe nucleus (Beier et al., 2015).

It was somewhat surprising to observe that dopamine loss did not affect the sucrose consumption in the fcHFHS paradigm. Indeed, anhedonic response towards sucrose solution has been clearly reported with manipulation of dopamine systems or loss of dopamine neurons (Der-Avakian and Markou, 2012; Faivre et al., 2020; Kaminska et al., 2017; Santiago et al., 2014; Zhang et al., 2016). However, dopamine deficient mice have higher rate of licking, bout size and duration with fewer total licks, without change in preference for sucrose (Cannon and Palmiter, 2003). Importantly, most of the anhedonia-related studies are carried with low to mild concentrations of sucrose solution (0.5% to 5%), while with higher concentrations of sucrose there can be a ceiling effect in the amount of sucrose taken daily (Wallace et al., 2008). It suggests that anhedonia-related studies are mostly highlighting dose-related shifts in sucrose preference rather than total loss of interest. In our study, we used the fcHFHS paradigm that relies on a high concentration of sucrose, which may mask dopamine-related modulation of the preference for sucrose (usually detected at thresholds doses) and explains why we did not observe any differences for overall sucrose consumption between 6-OHDA-lesioned and control animals.

With most NAc functional studies focusing on the mNAc, the importance of the lNAc has been mostly neglected. The present study provides behavioral evidence that dopamine in the lNAc is influential for food-type preference in food intake. Our results more specifically indicate that reducing the dopamine tone in this region was sufficient to favor fat intake in male rats. These findings provide a new insight into how dopamine influences food intake and food preference, opening the path for further research on lNAc regulation of fat intake in normal and pathological states.

## ABBREVIATIONS

6-OHDA: 6-hydroxydopamine;
ANOVA: analysis of variance;
AP: anteroposterior;
DA: dopamine;
DLS: dorsolateral striatum;
fcHFHS: free-choice high-fat high-sugar;
L: lateral;
lNAc: lateral nucleus accumbens;
mNAc: medial nucleus accumbens;
PB: phosphate buffer;
PBS: phosphate buffer saline;
PBS-T: PBS with Triton X-100;
SNc: substantia nigra pars compacta;
TH: tyrosine hydroxylase;
V: vertical;
VTA: ventral tegmental area

## AUTHOR CONTRIBUTIONS

A.J. designed and performed experiments, analyzed data, and drafted the manuscript; F.F. performed experiments. S.F. and M.B. supervised, designed the study, and drafted the manuscript.

## ACKNOWLEDGEMENTS

We thank the core facility Chronobiotron UMS3415 (Strasbourg, France) for animal care, and the UPS3156 *in vitro* imaging core facility (Strasbourg, France) for NanoZoomer scanning.

---

**Figure 1S:**
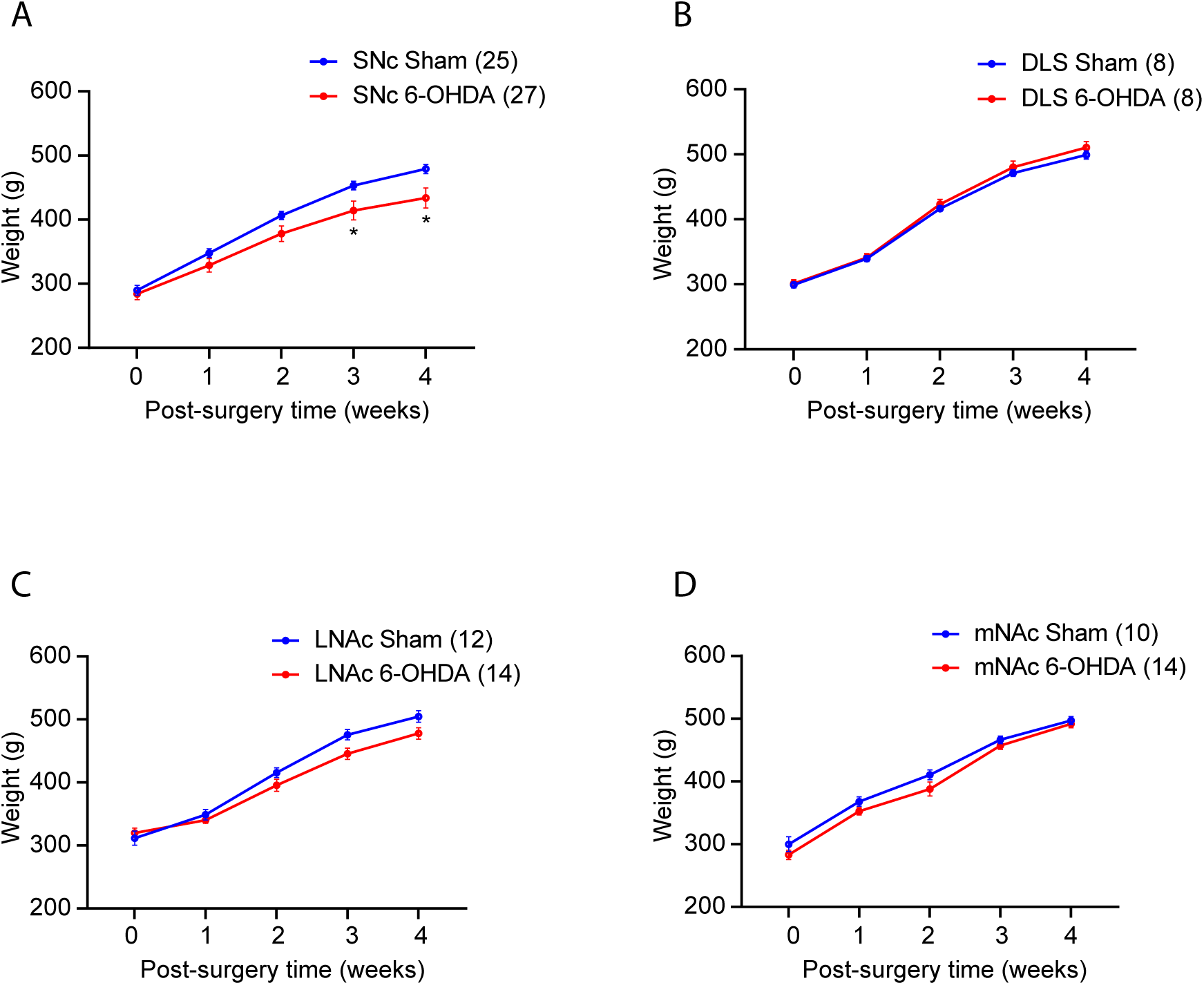
Rats’ bodyweight. Bodyweight was monitored throughout the experiments: (**A**) for SNc Sham and 6-OHDA lesioned animals, (**B**) for DLS Sham and 6-OHDA lesioned animals, (**C**) for lateral NAc Sham and 6-OHDA lesioned animals, and (**D**) for medial NAc Sham and 6-OHDA lesioned animals. The number of animals is given between brackets. Data are displayed as mean±SEM. *, p < 0.05. 6-OHDA, 6-hydroxydopamine; DLS, dorsolateral striatum; NAc, nucleus accumbens; SNc, substantia nigra pars compacta.

**Figure 2S:**
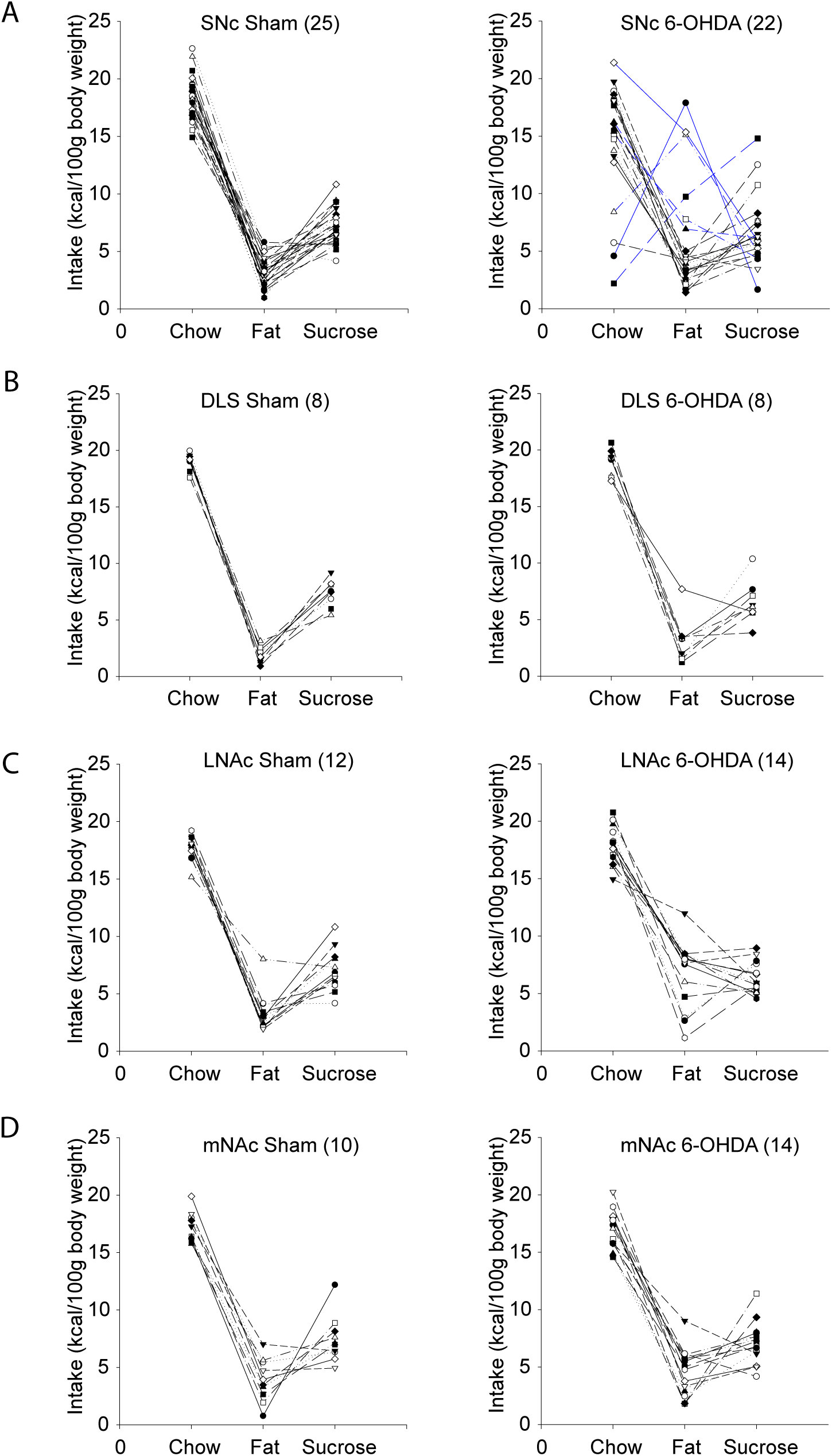
Individual patterns in the High-fat High-sugar free choice tests. The individual patterns are displayed as kcal intake per 100 g bodyweight for the three nutrient types for: (**A**) SNc Sham and 6-OHDA lesioned animals, (**B**) DLS Sham and 6-OHDA lesioned animals, (**C**) lateral NAc Sham and 6-OHDA lesioned animals, and (**D**) medial NAc Sham and 6-OHDA lesioned animals. The number of animals is given between brackets. 6-OHDA, 6-hydroxydopamine; DLS, dorsolateral striatum; NAc, nucleus accumbens; SNc, substantia nigra pars compacta.

## REFERENCES

Adams, W.K., Sussman, J.L., Kaur, S.,D’Souza A, M., Kieffer, T.J., and Winstanley, C.A. (2015). Long-term, calorie-restricted intake of a high-fat diet in rats reduces impulse control and ventral striatal D2 receptor signalling - two markers of addiction vulnerability. Eur J Neurosci 42, 3095–3104. doi:10.1111/ejn.13117

Alsio, J., Olszewski, P.K., Norback, A.H., Gunnarsson, Z.E., Levine, A.S., Pickering, C., and Schioth, H.B. (2010). Dopamine D1 receptor gene expression decreases in the nucleus accumbens upon long-term exposure to palatable food and differs depending on diet-induced obesity phenotype in rats. Neuroscience 171, 779–787. doi:10.1016/j.neuroscience.2010.09.046

Beier, K.T., Steinberg, E.E., DeLoach, K.E., Xie, S., Miyamichi, K., Schwarz, L., Gao, X.J., Kremer, E.J., Malenka, R.C., and Luo, L. (2015). Circuit Architecture of VTA Dopamine Neurons Revealed by Systematic Input-Output Mapping. Cell 162, 622–634. doi:10.1016/j.cell.2015.07.015

Berke, J.D. (2018). What does dopamine mean? Nat Neurosci 21, 787–793. doi:10.1038/s41593-018-0152-y

Bieri, G., Brahic, M., Bousset, L., Couthouis, J., Kramer, N.J., Ma, R., Nakayama, L., Monbureau, M., Defensor, E., Schule, B., et al. (2019). LRRK2 modifies alpha-syn pathology and spread in mouse models and human neurons. Acta Neuropathol 137, 961–980. doi:10.1007/s00401-019-01995-0

Boekhoudt, L., Roelofs, T.J.M., de Jong, J.W., de Leeuw, A.E., Luijendijk, M.C.M., Wolterink-Donselaar, I.G., van der Plasse, G., and Adan, R.A.H. (2017). Does activation of midbrain dopamine neurons promote or reduce feeding? Int J Obes (Lond) 41, 1131–1140. doi:10.1038/ijo.2017.74

Cannon, C.M., and Palmiter, R.D. (2003). Reward without dopamine. J Neurosci 23, 10827–10831. doi:10.1523/JNEUROSCI.23-34-10827.2003

Carr, K.D. (2020). Homeostatic regulation of reward via synaptic insertion of calcium-permeable AMPA receptors in nucleus accumbens. Physiol Behav 219, 112850. doi:10.1016/j.physbeh.2020.112850

Cooper, S.J., Francis, J., Al-Naser, H., and Barber, D. (1992). Evidence for dopamine D-1 receptor-mediated facilitatory and inhibitory effects on feeding behaviour in rats. J Psychopharmacol 6, 27–33. doi:10.1177/026988119200600108

Cox, J., and Witten, I.B. (2019). Striatal circuits for reward learning and decision-making. Nat Rev Neurosci 20, 482–494. doi:10.1038/s41583-019-0189-2

Cuvelier, E., Mequinion, M., Leghay, C., Sibran, W., Stievenard, A., Sarchione, A., Bonte, M.A., Vanbesien-Mailliot, C., Viltart, O., Saitoski, K., et al. (2018). Overexpression of Wild-Type Human Alpha-Synuclein Causes Metabolism Abnormalities in Thy1-aSYN Transgenic Mice. Front Mol Neurosci 11, 321. doi:10.3389/fnmol.2018.00321

da Silva, J.A., Tecuapetla, F., Paixao, V., and Costa, R.M. (2018). Dopamine neuron activity before action initiation gates and invigorates future movements. Nature 554, 244–248. doi:10.1038/nature25457

de Araujo, I.E., Oliveira-Maia, A.J., Sotnikova, T.D., Gainetdinov, R.R., Caron, M.G., Nicolelis, M.A., and Simon, S.A. (2008). Food reward in the absence of taste receptor signaling. Neuron 57, 930–941. doi:10.1016/j.neuron.2008.01.032

de Weijer, B.A., van de Giessen, E., van Amelsvoort, T.A., Boot, E., Braak, B., Janssen, I.M., van de Laar, A., Fliers, E., Serlie, M.J., and Booij, J. (2011). Lower striatal dopamine D2/3 receptor availability in obese compared with non-obese subjects. EJNMMI Res 1, 37. doi:10.1186/2191-219X-1-37

Der-Avakian, A., and Markou, A. (2012). The neurobiology of anhedonia and other reward-related deficits. Trends Neurosci 35, 68–77. doi:10.1016/j.tins.2011.11.005

DiFeliceantonio, A.G., Coppin, G., Rigoux, L., Edwin Thanarajah, S., Dagher, A., Tittgemeyer, M., and Small, D.M. (2018). Supra-Additive Effects of Combining Fat and Carbohydrate on Food Reward. Cell Metab 28, 33–44 e33. doi:10.1016/j.cmet.2018.05.018

Durst, M., Konczol, K., Balazsa, T., Eyre, M.D., and Toth, Z.E. (2019). Reward-representing D1-type neurons in the medial shell of the accumbens nucleus regulate palatable food intake. Int J Obes (Lond) 43, 917–927. doi:10.1038/s41366-018-0133-y

Faivre, F., Joshi, A., Bezard, E., and Barrot, M. (2019). The hidden side of Parkinson’s disease: Studying pain, anxiety and depression in animal models. Neurosci Biobehav Rev 96, 335–352. doi:10.1016/j.neubiorev.2018.10.004

Faivre, F., Sanchez-Catalan, M.J., Dovero, S., Bido, S., Joshi, A., Bezard, E., and Barrot, M. (2020). Ablation of the tail of the ventral tegmental area compensates symptoms in an experimental model of Parkinson’s disease. Neurobiol Dis 139, 104818. doi:10.1016/j.nbd.2020.104818

Fernandes, A.B., Alves da Silva, J., Almeida, J., Cui, G., Gerfen, C.R., Costa, R.M., and Oliveira-Maia, A.J. (2020). Postingestive Modulation of Food Seeking Depends on Vagus-Mediated Dopamine Neuron Activity. Neuron 106, 1–11. doi:10.1016/j.neuron.2020.03.009

Ferreira, J.G., Tellez, L.A., Ren, X., Yeckel, C.W., and de Araujo, I.E. (2012). Regulation of fat intake in the absence of flavour signalling. J Physiol 590, 953–972. doi:10.1113/jphysiol.2011.218289

Fletcher, P.C., and Kenny, P.J. (2018). Food addiction: a valid concept? Neuropsychopharmacology 43, 2506–2513. doi:10.1038/s41386-018-0203-9

Fritz, B.M., Munoz, B., Yin, F., Bauchle, C., and Atwood, B.K. (2018). A High-fat, High-sugar ‘Western’ Diet Alters Dorsal Striatal Glutamate, Opioid, and Dopamine Transmission in Mice. Neuroscience 372, 1–15. doi:10.1016/j.neuroscience.2017.12.036

Gerfen, C.R., and Bolam, J.P. (2016). Chapter 1 - The Neuroanatomical Organization of the Basal Ganglia. In Handbook of Behavioral Neuroscience. H. Steiner, and K.Y. Tseng, eds. (Elsevier), pp. 3–32. doi:10.1016/B978-0-12-802206-1.00001-5

Giasson, B.I., Duda, J.E., Quinn, S.M., Zhang, B., Trojanowski, J.Q., and Lee, V.M. (2002). Neuronal alpha-synucleinopathy with severe movement disorder in mice expressing A53T human alpha-synuclein. Neuron 34, 521–533. doi:10.1016/s0896-6273(02)00682-7

Gispert, S., Ricciardi, F., Kurz, A., Azizov, M., Hoepken, H.H., Becker, D., Voos, W., Leuner, K., Muller, W.E., Kudin, A.P., et al. (2009). Parkinson phenotype in aged PINK1-deficient mice is accompanied by progressive mitochondrial dysfunction in absence of neurodegeneration. PLoS One 4, e5777. doi:10.1371/journal.pone.0005777

Haber, S.N., Fudge, J.L., and McFarland, N.R. (2000). Striatonigrostriatal pathways in primates form an ascending spiral from the shell to the dorsolateral striatum. J Neurosci 20, 2369–2382. doi:10.1523/JNEUROSCI.20-06-02369.2000

Hajnal, A., Smith, G.P., and Norgren, R. (2004). Oral sucrose stimulation increases accumbens dopamine in the rat. Am J Physiol Regul Integr Comp Physiol 286, R31–37. doi:10.1152/ajpregu.00282.2003

Hamid, A.A., Pettibone, J.R., Mabrouk, O.S., Hetrick, V.L., Schmidt, R., Vander Weele, C.M., Kennedy, R.T., Aragona, B.J., and Berke, J.D. (2016). Mesolimbic dopamine signals the value of work. Nat Neurosci 19, 117–126. doi:10.1038/nn.4173

Han, W., Tellez, L.A., Perkins, M.H., Perez, I.O., Qu, T., Ferreira, J., Ferreira, T.L., Quinn, D., Liu, Z.W., Gao, X.B., et al. (2018). A Neural Circuit for Gut-Induced Reward. Cell 175, 665–678 e623. doi:10.1016/j.cell.2018.08.049

Hernandez, L., and Hoebel, B.G. (1988). Feeding and hypothalamic stimulation increase dopamine turnover in the accumbens. Physiol Behav 44, 599–606. doi:10.1016/0031-9384(88)90324-1

Hnasko, T.S., Perez, F.A., Scouras, A.D., Stoll, E.A., Gale, S.D., Luquet, S., Phillips, P.E., Kremer, E.J., and Palmiter, R.D. (2006). Cre recombinase-mediated restoration of nigrostriatal dopamine in dopamine-deficient mice reverses hypophagia and bradykinesia. Proc Natl Acad Sci U S A 103, 8858–8863. doi:10.1073/pnas.0603081103

Hoebel, B.G. (1985). Brain neurotransmitters in food and drug reward. Am J Clin Nutr 42, 1133–1150. doi:10.1093/ajcn/42.5.1133

Howe, M.W., and Dombeck, D.A. (2016). Rapid signalling in distinct dopaminergic axons during locomotion and reward. Nature 535, 505–510. doi:10.1038/nature18942

Hryhorczuk, C., Florea, M., Rodaros, D., Poirier, I., Daneault, C., Des Rosiers, C., Arvanitogiannis, A., Alquier, T., and Fulton, S. (2016). Dampened Mesolimbic Dopamine Function and Signaling by Saturated but not Monounsaturated Dietary Lipids. Neuropsychopharmacology 41, 811–821. doi:10.1038/npp.2015.207

Hu, H. (2016). Reward and Aversion. Annu Rev Neurosci 39, 297–324. doi:10.1146/annurev-neuro-070815-014106

Huang, X.F., Yu, Y., Zavitsanou, K., Han, M., and Storlien, L. (2005). Differential expression of dopamine D2 and D4 receptor and tyrosine hydroxylase mRNA in mice prone, or resistant, to chronic high-fat diet-induced obesity. Brain Res Mol Brain Res 135, 150–161. doi:10.1016/j.molbrainres.2004.12.013

Huang, X.F., Zavitsanou, K., Huang, X., Yu, Y., Wang, H., Chen, F., Lawrence, A.J., and Deng, C. (2006). Dopamine transporter and D2 receptor binding densities in mice prone or resistant to chronic high fat diet-induced obesity. Behav Brain Res 175, 415–419. doi:10.1016/j.bbr.2006.08.034

Ilango, A., Kesner, A.J., Keller, K.L., Stuber, G.D., Bonci, A., and Ikemoto, S. (2014). Similar roles of substantia nigra and ventral tegmental dopamine neurons in reward and aversion. J Neurosci 34, 817–822. doi:10.1523/JNEUROSCI.1703-13.2014

Johnson, P.M., and Kenny, P.J. (2010). Dopamine D2 receptors in addiction-like reward dysfunction and compulsive eating in obese rats. Nat Neurosci 13, 635–641. doi:10.1038/nn.2519

Kaminska, K., Lenda, T., Konieczny, J., Czarnecka, A., and Lorenc-Koci, E. (2017). Depressive-like neurochemical and behavioral markers of Parkinson’s disease after 6-OHDA administered unilaterally to the rat medial forebrain bundle. Pharmacol Rep 69, 985–994. doi:10.1016/j.pharep.2017.05.016

Kistner, A., Lhommee, E., and Krack, P. (2014). Mechanisms of body weight fluctuations in Parkinson’s disease. Front Neurol 5, 84. doi:10.3389/fneur.2014.00084

Koob, G.F., Riley, S.J., Smith, S.C., and Robbins, T.W. (1978). Effects of 6-hydroxydopamine lesions of the nucleus accumbens septi and olfactory tubercle on feeding, locomotor activity, and amphetamine anorexia in the rat. J Comp Physiol Psychol 92, 917–927. doi:10.1037/h0077542

la Fleur, S.E., Luijendijk, M.C., van der Zwaal, E.M., Brans, M.A., and Adan, R.A. (2014). The snacking rat as model of human obesity: effects of a free-choice high-fat high-sugar diet on meal patterns. Int J Obes (Lond) 38, 643–649. doi:10.1038/ijo.2013.159

la Fleur, S.E., Luijendijk, M.C., van Rozen, A.J., Kalsbeek, A., and Adan, R.A. (2011). A free-choice high-fat high-sugar diet induces glucose intolerance and insulin unresponsiveness to a glucose load not explained by obesity. Int J Obes (Lond) 35, 595–604. doi:10.1038/ijo.2010.164

la Fleur, S.E., van Rozen, A.J., Luijendijk, M.C., Groeneweg, F., and Adan, R.A. (2010). A free-choice high-fat high-sugar diet induces changes in arcuate neuropeptide expression that support hyperphagia. Int J Obes (Lond) 34, 537–546. doi:10.1038/ijo.2009.257

Lammel, S., Lim, B.K., Ran, C., Huang, K.W., Betley, M.J., Tye, K.M., Deisseroth, K., and Malenka, R.C. (2012). Input-specific control of reward and aversion in the ventral tegmental area. Nature 491, 212–217. doi:10.1038/nature11527

Lardeux, S., Kim, J.J., and Nicola, S.M. (2015). Intermittent-access binge consumption of sweet high-fat liquid does not require opioid or dopamine receptors in the nucleus accumbens. Behav Brain Res 292, 194–208. doi:10.1016/j.bbr.2015.06.015

Lee, K., Claar, L.D., Hachisuka, A., Bakhurin, K.I., Nguyen, J., Trott, J.M., Gill, J.L., and Masmanidis, S.C. (2020). Temporally restricted dopaminergic control of reward-conditioned movements. Nat Neurosci 23, 209–216. doi:10.1038/s41593-019-0567-0

Li, X., Redus, L., Chen, C., Martinez, P.A., Strong, R., Li, S., and O’Connor, J.C. (2013). Cognitive dysfunction precedes the onset of motor symptoms in the MitoPark mouse model of Parkinson’s disease. PLoS One 8, e71341. doi:10.1371/journal.pone.0071341

Liang, N.C., Hajnal, A., and Norgren, R. (2006). Sham feeding corn oil increases accumbens dopamine in the rat. Am J Physiol Regul Integr Comp Physiol 291, R1236–1239. doi:10.1152/ajpregu.00226.2006

London, T.D., Licholai, J.A., Szczot, I., Ali, M.A., LeBlanc, K.H., Fobbs, W.C., and Kravitz, A.V. (2018). Coordinated Ramping of Dorsal Striatal Pathways preceding Food Approach and Consumption. J Neurosci 38, 3547–3558. doi:10.1523/JNEUROSCI.2693-17.2018

Ma, K., Xiong, N., Shen, Y., Han, C., Liu, L., Zhang, G., Wang, L., Guo, S., Guo, X., Xia, Y., et al. (2018). Weight Loss and Malnutrition in Patients with Parkinson’s Disease: Current Knowledge and Future Prospects. Front Aging Neurosci 10, 1. doi:10.3389/fnagi.2018.00001

Martin-Iverson, M.T., and Dourish, C.T. (1988). Role of dopamine D-1 and D-2 receptor subtypes in mediating dopamine agonist effects on food consumption in rats. Psychopharmacology (Berl) 96, 370–374. doi:10.1007/BF00216064

Morales, M., and Margolis, E.B. (2017). Ventral tegmental area: cellular heterogeneity, connectivity and behaviour. Nat Rev Neurosci 18, 73–85. doi:10.1038/nrn.2016.165

Munoz-Manchado, A.B., Villadiego, J., Romo-Madero, S., Suarez-Luna, N., Bermejo-Navas, A., Rodriguez-Gomez, J.A., Garrido-Gil, P., Labandeira-Garcia, J.L., Echevarria, M., Lopez-Barneo, J., et al. (2016). Chronic and progressive Parkinson’s disease MPTP model in adult and aged mice. J Neurochem 136, 373–387. doi:10.1111/jnc.13409

O’Connor, E.C., Kremer, Y., Lefort, S., Harada, M., Pascoli, V., Rohner, C., and Luscher, C. (2015). Accumbal D1R Neurons Projecting to Lateral Hypothalamus Authorize Feeding. Neuron 88, 553–564. doi:10.1016/j.neuron.2015.09.038

Palmiter, R.D. (2008). Dopamine signaling in the dorsal striatum is essential for motivated behaviors: lessons from dopamine-deficient mice. Ann N Y Acad Sci 1129, 35–46. doi:10.1196/annals.1417.003

Paxinos, G., and Watson, C. (2013). The Rat Brain in Stereotaxic Coordinates. (Academic Press).

Porras, G., Li, Q., and Bezard, E. (2012). Modeling Parkinson’s disease in primates: The MPTP model. Cold Spring Harb Perspect Med 2, a009308. doi:10.1101/cshperspect.a009308

Rada, P., Avena, N.M., Barson, J.R., Hoebel, B.G., and Leibowitz, S.F. (2012). A high-fat meal, or intraperitoneal administration of a fat emulsion, increases extracellular dopamine in the nucleus accumbens. Brain Sci 2, 242–253. doi:10.3390/brainsci2020242

Santiago, R.M., Barbiero, J., Gradowski, R.W., Bochen, S., Lima, M.M., Da Cunha, C., Andreatini, R., and Vital, M.A. (2014). Induction of depressive-like behavior by intranigral 6-OHDA is directly correlated with deficits in striatal dopamine and hippocampal serotonin. Behav Brain Res 259, 70–77. doi:10.1016/j.bbr.2013.10.035

Sharma, S., and Fulton, S. (2013). Diet-induced obesity promotes depressive-like behaviour that is associated with neural adaptations in brain reward circuitry. Int J Obes (Lond) 37, 382–389. doi:10.1038/ijo.2012.48

Sotak, B.N., Hnasko, T.S., Robinson, S., Kremer, E.J., and Palmiter, R.D. (2005). Dysregulation of dopamine signaling in the dorsal striatum inhibits feeding. Brain Res 1061, 88–96. doi:10.1016/j.brainres.2005.08.053

South, T., and Huang, X.F. (2008). High-fat diet exposure increases dopamine D2 receptor and decreases dopamine transporter receptor binding density in the nucleus accumbens and caudate putamen of mice. Neurochem Res 33, 598–605. doi:10.1007/s11064-007-9483-x

Sundstrom, E., Fredriksson, A., and Archer, T. (1990). Chronic neurochemical and behavioral changes in MPTP-lesioned C57BL/6 mice: a model for Parkinson’s disease. Brain Res 528, 181–188. doi:10.1016/0006-8993(90)91656-2

Szczypka, M.S., Kwok, K., Brot, M.D., Marck, B.T., Matsumoto, A.M., Donahue, B.A., and Palmiter, R.D. (2001). Dopamine production in the caudate putamen restores feeding in dopamine-deficient mice. Neuron 30, 819–828. doi:10.1016/s0896-6273(01)00319-1

Tellez, L.A., Medina, S., Han, W., Ferreira, J.G., Licona-Limon, P., Ren, X., Lam, T.T., Schwartz, G.J., and de Araujo, I.E. (2013). A gut lipid messenger links excess dietary fat to dopamine deficiency. Science 341, 800–802. doi:10.1126/science.1239275

Tsai, H.C., Zhang, F., Adamantidis, A., Stuber, G.D., Bonci, A., de Lecea, L., and Deisseroth, K. (2009). Phasic firing in dopaminergic neurons is sufficient for behavioral conditioning. Science 324, 1080–1084. doi:10.1126/science.1168878

Ungerstedt, U. (1968). 6-Hydroxy-dopamine induced degeneration of central monoamine neurons. Eur J Pharmacol 5, 107–110. doi:10.1016/0014-2999(68)90164-7

Ungerstedt, U. (1971). Adipsia and aphagia after 6-hydroxydopamine induced degeneration of the nigro-striatal dopamine system. Acta Physiol Scand Suppl 367, 95–122. doi:10.1111/j.1365-201x.1971.tb11001.x

van de Giessen, E., la Fleur, S.E., Eggels, L., de Bruin, K., van den Brink, W., and Booij, J. (2013). High fat/carbohydrate ratio but not total energy intake induces lower striatal dopamine D2/3 receptor availability in diet-induced obesity. Int J Obes (Lond) 37, 754–757. doi:10.1038/ijo.2012.128

Volkow, N.D., Wise, R.A., and Baler, R. (2017). The dopamine motive system: implications for drug and food addiction. Nat Rev Neurosci 18, 741–752. doi:10.1038/nrn.2017.130

Wallace, D.L., Vialou, V., Rios, L., Carle-Florence, T.L., Chakravarty, S., Kumar, A., Graham, D.L., Green, T.A., Kirk, A., Iniguez, S.D., et al. (2008). The influence of DeltaFosB in the nucleus accumbens on natural reward-related behavior. J Neurosci 28, 10272–10277. doi:10.1523/JNEUROSCI.1531-08.2008

Wang, G.J., Volkow, N.D., Logan, J., Pappas, N.R., Wong, C.T., Zhu, W., Netusil, N., and Fowler, J.S. (2001). Brain dopamine and obesity. Lancet 357, 354–357. doi:10.1016/s0140-6736(00)03643-6

Witten, I.B., Steinberg, E.E., Lee, S.Y., Davidson, T.J., Zalocusky, K.A., Brodsky, M., Yizhar, O., Cho, S.L., Gong, S., Ramakrishnan, C., et al. (2011). Recombinase-driver rat lines: tools, techniques, and optogenetic application to dopamine-mediated reinforcement. Neuron 72, 721–733. doi:10.1016/j.neuron.2011.10.028

Yang, H., de Jong, J.W., Tak, Y., Peck, J., Bateup, H.S., and Lammel, S. (2018). Nucleus Accumbens Subnuclei Regulate Motivated Behavior via Direct Inhibition and Disinhibition of VTA Dopamine Subpopulations. Neuron 97, 434–449 e434. doi:10.1016/j.neuron.2017.12.022

Zhang, Q.S., Heng, Y., Mou, Z., Huang, J.Y., Yuan, Y.H., and Chen, N.H. (2017). Reassessment of subacute MPTP-treated mice as animal model of Parkinson’s disease. Acta Pharmacol Sin 38, 1317–1328. doi:10.1038/aps.2017.49

Zhang, Y.M., Zhang, L., Wang, Y., Sun, Y.N., Guo, Y., Du, C.X., Zhang, J., Yao, L., Yu, S.Q., and Liu, J. (2016). Activation and blockade of prelimbic 5-HT6 receptors produce different effects on depressive-like behaviors in unilateral 6-hydroxydopamine-induced Parkinson’s rats. Neuropharmacology 110, 25–36. doi:10.1016/j.neuropharm.2016.07.014

Zhou, Q.Y., and Palmiter, R.D. (1995). Dopamine-deficient mice are severely hypoactive, adipsic, and aphagic. Cell 83, 1197–1209. doi:10.1016/0092-8674(95)90145-0

Zimmerman, C.A., and Knight, Z.A. (2020). Layers of signals that regulate appetite. Curr Opin Neurobiol 64, 79–88. doi:10.1016/j.conb.2020.03.007

